# Artificial reefs geographical location matters more than shape, age and depth for sessile invertebrate colonization in the Gulf of Lion (NorthWestern Mediterranean Sea)

**DOI:** 10.1101/2021.10.08.463669

**Authors:** Sylvain Blouet, Lorenzo Bramanti, Katell Guizien

## Abstract

Artificial reefs (ARs) have been used to support fishing activities. Sessile invertebrates are essential components of trophic networks within ARs, supporting fish productivity. However, colonization by sessile invertebrates is possible only after effective larval dispersal from source populations, usually in natural habitat. While most studies focused on short term colonization by pioneer species, we propose to test the relevance of geographic location, shape, age and depth of immersion on the ARs long term colonization by species found in natural stable communities in the Gulf of Lion. We recorded the presence of five sessile invertebrates species, with contrasting life history traits and regional distribution in the natural rocky habitat, on ARs with different shapes deployed during two immersion time periods (1985 and the 2000s) and in two depth ranges (<20m and >20m). At the local level (∼5kms), neither shape, depth nor immersion duration differentiated ARs assemblages. At the regional scale (>30kms), colonization patterns differed between species, resulting in diverse assemblages. This study highlights the primacy of geographical positioning over shape, immersion duration and depth in ARs colonization, suggesting it should be accounted for in maritime spatial planning.

## Introduction

The decline of fish stocks and natural marine habitat degradation resulting from human exploitation have been documented worldwide for decades (Claudet and Fraschetti, 2010; Jackson, 2001; Pauly et al., 2002).

Artificial reefs (ARs) have been primarily implemented to reduce the pressure of fisheries in coastal areas, complementing other management tools such as marine protected areas or regulatory measures such as fishing licenses (Claudet and Pelletier, 2004; Seaman, 2007; Wilson, 2002). Moreover, ARs could provide economic benefits such as recreational and traditional fishing and scuba diving (Chen et al., 2013). Beneficial effects such as increase in fish biomass and capture efficiency near ARs have been reported (reviewed by Bohnsack and Sutherland, 1985; Tessier et al., 2014) but led to a debate on the effects of ARs fishery, opposing attraction vs production (Grossman et al., 1997). The fish attraction argument is based on the quick colonization by fishes and mobile invertebrates (Powers et al., 2003; Relini, 2002; Santos and Monteiro, 2007). The fish production argument is based on the hypotheses of a better protection against predators and an increase in available substrate area for larval establishment thanks to habitat complexification and an increase of available trophic resource (Bohnsack, 1989). In natural rocky habitats, benthic invertebrates play an essential role in fish trophic networks (Ardizzone et al., 1996; Martens et al., 2006). ARs trophic network showed similarity with natural rocky habitat, with dominance of filter-feeders using phytoplanktonic primary production and fish predation on crustacean colonizing the ARs (Cresson, 2013). Moreover, AR deployed in sandy areas are expected to enhance fish productivity given that epifauna secondary production per ARs unit surface has been estimated to be 30 times greater than that of natural sandy infauna (Steimle, 2002). However, those estimates were made shortly after immersion and do not prove the long-term fish production in ARs and supporting this argument would require extending data on colonization by benthic invertebrates in the long-term (Svane and Petersen, 2001). Indeed, age since deployment has been described as a key factor to explain ARs coverage by benthic invertebrates (Svane and Petersen, 2001). The assemblages of benthic communities are expected to change over time in a succession between pioneer and specialist species (Connell and Slatyer, 1977). In contrast with pioneer species, specialist ones have slower colonization dynamics, because of their lower fecundity (Fava et al., 2016). However, after colonization, specialists are expected to outcompete pioneer species due to their more efficient use of environmental resources (Connell and Slatyer, 1977). Among these, light availability is an essential factor shaping marine benthic communities across the water depth gradient (Odum, 1971). Several studies have shown a decrease in the density of benthic invertebrates with depth on ARs (Lewbel et al., 1986; Moura et al., 2007; Shinn and Wicklund, 1989; Van der Stap et al., 2016) explained by the decrease in light intensity (Relini et al., 1994). The structural complexity has also been put forward as important characteristics linked to ARs efficiency in ecological restoration (Strain et al., 2018). Structural complexity increases available surface for colonization and niches diversity with various shelter and light exposure conditions, the latter being related to different benthic assemblage compositions (Glasby, 2000; Glasby, 1999) and higher productivity (Vivier et al., 2021). The recent 3D printing techniques using concrete allow the design of ARs mimicking natural habitats (Ly et al., 2021). However, those studies concerned short-term colonization (<3.5 years) (Wendt et al., 1989) hence based on pioneer species with high dispersal capacities which long term colonization is likely regulated by post-settlement processes such as competition, predation and physical disturbance (Todd, 1998). In contrast to mobile species, sessile benthic invertebrates can only colonize reefs after larval dispersal which is limited by reproduction frequency (Thorson, 1950). Colonization implies thus an effective dispersal between natural areas and ARs, which depends on source population spatial distribution, species fecundity, dispersive larval traits and ocean circulation. Nevertheless, until now dispersal drivers have been disregarded while colonization disparities among ARs may result from differences in both larval connectivity (which in turn depends on fecundity, dispersal capacities and adult distribution in the natural habitat) and post-recruitment processes.

The objective of the present study was to test the hypothesis that the geographical location of ARs deployment with respect to the natural habitat can condition ARs colonization after more than 10 years. To this aim we investigated the effects of local (shape, depth and immersion duration) and regional (geographic area) factors on the presence of five species of sessile invertebrates with different life history traits, endemics to the Gulf of Lion (GoL) (Northwestern Mediterranean Sea) and frequently found on natural hard substrates.

## Methods

### Study area and spatially stratified sampling design

The study area extended along 160 km of the GoL coastline (Figure 1). The GoL is a wide microtidal continental shelf dominated by soft-bottom habitat with few small rocky habitat patches of less than 20 km^2^. The GoL is a homogeneous and isolated hydrodynamic unit (Rossi et al., 2014), delimited by the northern current (Millot, 1990).

**Figure 1.**
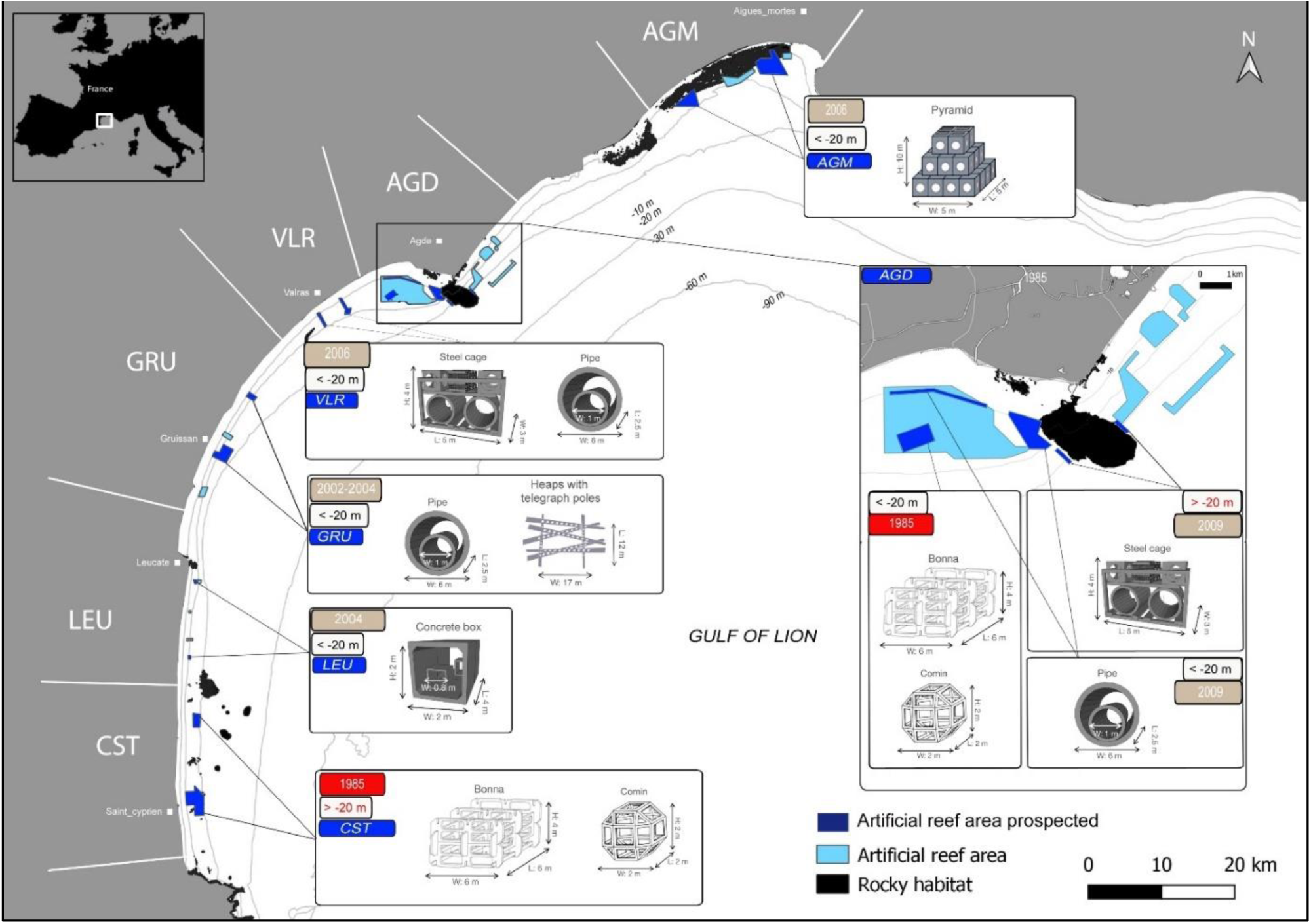
Map showing the layout of the 6 geographical sectors (AGM, AGD, VLR, GRU, LEU, CST) and the 16 sites prospected (indicated in dark blue) for artificial reef sampling. Areas indicated in light blue corresponded to artificial reef deployment areas. Areas indicated in black corresponded to natural hard-substrate areas. Some of the AR shape illustrations were taken from Tessier et al., 2014.

Between 1985 and 2009, 763 ARs with different shapes or material for a total volume of 37 575 m^3^ (Tessier et al., 2015) have been deployed along the GoL coastline over 66 km² of state concessions (Cepralmar, 2015) between 10 and 35 m depth (Figure 1, Blouet et al., 2021). The ARs deployed in GoL represent 40% of the total AR volume in France (Tessier et al., 2015).

In the present study we examined the oldest (in the 80s) and the youngest (in the 2000s) ARs with the most common shapes (namely pipe, steel cage, Bonna, Comin, pyramid, heap of telegraph poles, and concrete box, Figure 1). For the analysis of data, we followed a stratified sampling design. To this aim, the GoL coastline was regularly divided into 6 geographical sectors separated by a distance ranging from 12 to 117 km, with a median value of 49 km (AGM referring to Aigues-Mortes, AGD to Agde, VLR to Valras, GRU to Gruissan, LEU to Leucate, CST to Canet/Saint-Cyprien sectors, Figure 1). Each sector included ARs deployed during either the first (1985, CST geographical sector), the last (2002-2009, AGM, VLR, GRU, LEU geographical sectors) or both deployment periods (AGD), and of different shapes. In each geographical sector, two sites were defined except in AGD (5 sites, AGD1, AGD2, AGD3, AGD4, AGD5) and GRU (3 sites, GRU1, GRU2, GRU3). To test for location effect, sites were separated by a distance ranging from 2.1 to 11.7 km (median value of 7.5 km) except in GRU, where two of the three sites (GRU1 and GRU2) were defined in the same location to account for different shapes. Hence, depending on the geographical sector, sites may differ either by AR shape, immersion depth or immersion duration (Figure 1). Due to this set up, the effect of these three factors was only tested locally (see Statistical analysis). In each site (except in the geographical sector GRU), three sampling units separated by a distance between 4 m and 3.6 km (median distance of 251 m) were set out by pooling neighboring ARs reaching a minimum surface of 89 m^2^ per sampling unit and totalling a minimum developed surface of 306 m^2^ per site (Table 1: Supplementary Material). In the geographical sector GRU, GRU2 and GRU3 included only one sampling unit because the surface of a single AR in these sites already yielded 459 m^2^ (heap of telegraph poles; Table 1: Supplementary Material). Such large continuous sampling units in each site aimed at limiting the effect of recruitment spatial variability over distances from 100s meters to kilometers arising from the nonuniformity of the flow of larvae (Daigle et al., 2014; Glasby, 2000; Simpson et al., 2017; Smale, 2012). Such a spatial scale is consistent with the spatial scale of flow homogeneity obtained in simulations over GoL soft-bottom habitat (Briton et al., 2018). This inventory methodology enabled us to test for the existence of structuring factors at the local and regional scales. Developed reef surface was calculated on the basis of technical specifications data present in the state concession documents taking into account only the colonizable surface (surfaces in contact with the sediment were excluded).

**Table 1.** Life history traits (spawning period, relative fecundity, life expectancy, age at sexual maturity, larval type, planktonic larval duration (PLD), relative abundance in the GoL) for the five species inventoried on ARs.

| Species | Spawning period | Fecundity | Life expectancy | Age at sexual maturity | Larval type | Planktonic larval duration (PLD) | Abundance in the GoL |
| --- | --- | --- | --- | --- | --- | --- | --- |
| *Leptogorgia sarmentosa* | June - August | ** | 20 years | 2-3 years | Lecithotrophic | 1-4 weeks (estimated from other gorgonians) | ** |
| *Eunicella singularis* | July - August | * | 25-30 years | 6 years | Lecithotrophic | 1-4 weeks (in aquarium) | *** |
| *Pentapora fascialis* | June | ** | 10 years | 2 years | Lecithotrophic | <1 days | ** |
| *Sabella spallanzani* | January - February | *** | 5 years | 1 years | Lecithotrophic | 21 days (in aquarium) | * |
| *Halocynthia papillosa* | September - October | *** (estimated from other ascidians) | unknown | 2 months (estimated from other ascidians) | lecithotrophic (estimated from other ascidians) | <12 hours (estimated from other ascidians) | ** |

Symbol legend: \* = low, \*\*=medium \*\*\*=strong

### Colonization assessment and species selection

Assessment of ARs colonization was carried out by autonomous scuba-diving in 2020 by direct visual census counting the number of individuals of the five target species on the entire surface of ARs (7295 m²; Table1: Supplementary Material). On the ARs, all individuals older than one year (size > 2 cm) have been recorded. Among the species listed in previous ARs inventories in the GoL, we selected five species that were present in most inventories, easy to identify by direct visual census and spanning different phyla with contrasting life-history traits (Créocean, 2003 & 2004; Table 1). We selected two gorgonians *Eunicella singularis* (Esper, 1791) and *Leptogorgia sarmentosa* (Esper, 1789), one bryozoan *Pentapora fascialis* (Pallas, 1766), one annelida *Sabella spallanzanii* (Gmelin, 1791) and one ascidian *Halocynthia papillosa* (Linnaeus, 1797) (Figure 2). The five species have a similar wide natural repartition area along European coasts ranging from 1 m to 250 m depth (Giangrande et al., 2005; Gori et al., 2011; Ponti et al., 2019; Turon, 1990; Weinberg and Weinberg, 1979). In addition, *S. spallanzanii* has been recorded along the coasts of Brazil, Australia and New Zealand where it is classified as an invasive nonindigenous species (Currie et al., 2000). The five species are present in the rocky habitat of the NW Mediterranean Sea (Laubier, 1966; True, 1970; Hong, 1980). However, in the GoL, where natural rocky habitat covers uneven surfaces within the 6 geographical sectors (from 3,123 10^7^ m^2^ for the AGM sector to 5 10^5^ m^2^ for the LEU sector), the five species display different spatial distributions (Dutrieux et al., 2005; Dalias et al., 2011; Guizien et al., 2022; S. Blouet personal observation). *E. singularis* is frequently observed and abundant throughout the GoL (from the AGM sector to the CST sector), while *L. sarmentosa*, less abundant, is present mainly in the center of the GoL (AGD, VLR, LEU, CST). *P. fascialis* is abundant in the west of the GoL (AGD, LEU, CST, and south of CST). The distribution of *H. papillosa* is not well known, however the species has been observed in all the rocky areas of the GoL. *S. spallanzanii* is present but rare in natural rocky habitat. Nevertheless, *S. spallanzanii* is very abundant in lagoons, ports and marinas of the GoL (S. Blouet personal observation) which has been indicated as preferred habitat of the species (Currie et al., 2000).

**Figure 2.**
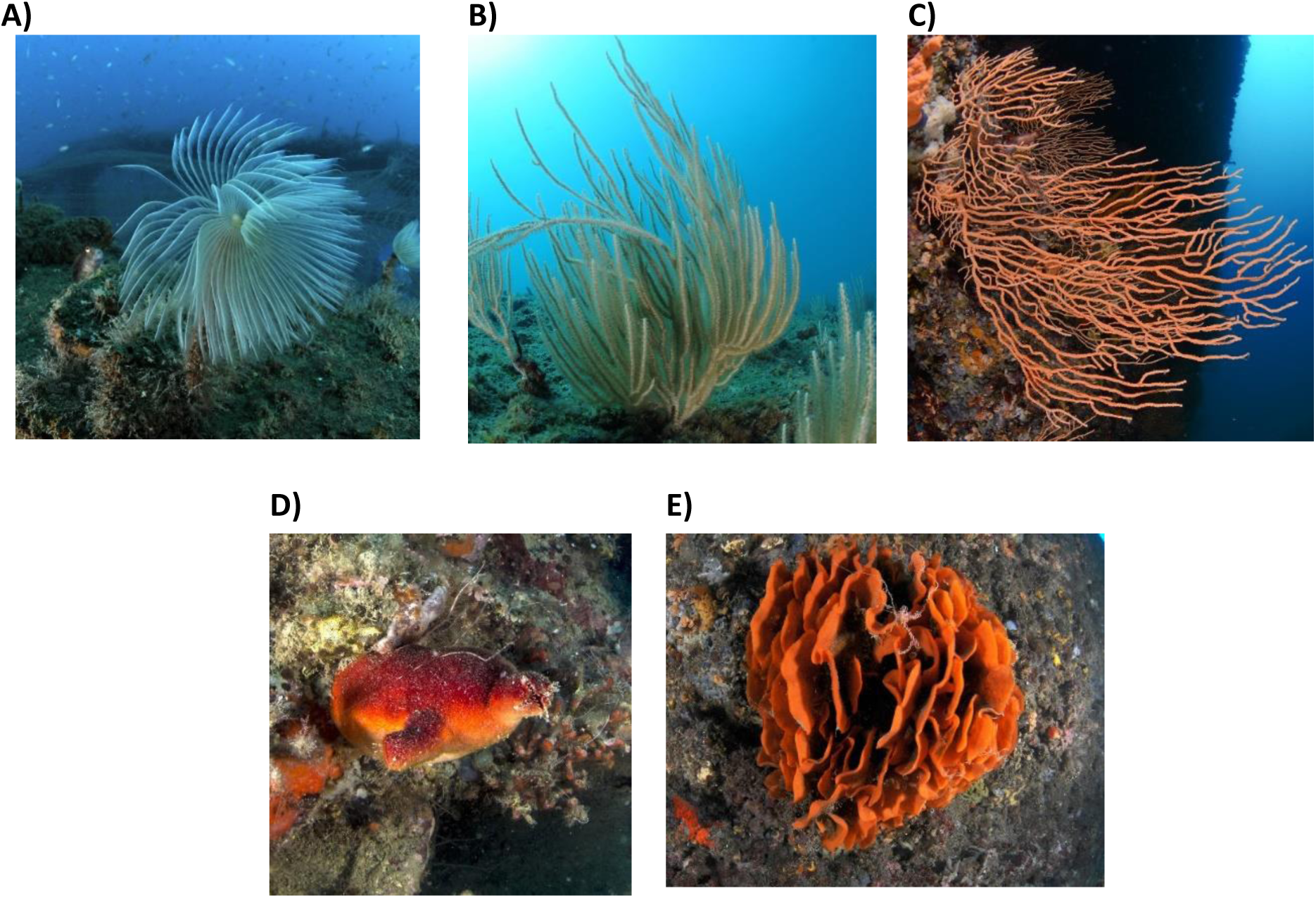
Photographs of the five species inventoried on AR *a) Sabella spallanzanii, b) Eunicella singularis,c) Leptogorgia sarmentosa, d) Halocynthia papillosa and* e) *Pentapora fascialis; all © Blouet sylvain*

The five species display different life-history traits. All five species reproduce once a year in different seasons and with different strategies (Table 1).

*S. spallanzani* reproduces in January-February, when water temperature is the coldest. The species displays multiple reproductive strategies: internal fertilization, with larvae brooded either inside or outside the mineral tube secreted around the body, and external fertilization broadcast spawning (Giangrande et al., 2000). In addition, asexual reproduction by fission has been reported (Read et al., 2011). *S. spallanzani* releases lecithotrophic larvae with a planktonic larval duration (PLD) of about 4 weeks (Giangrande et al., 2000). Its life span can exceed 5 years, with sexual maturity after one year (Giangrande and Petraroli 1994; Giangrande et al., 2000).

Like most gorgonians, *E. singularis* releases lecithotrophic larvae in early summer (June to August). Even though larval competency period can reach up to 2 months (Guizien et al., 2020; Zelli et al., 2020), PLDs ranging from 7 to 14 days best explained gene flow among *E. singularis* natural populations dwelling in the fragmented rocky habitat of the GoL (Padron et al., 2018). *E. singularis* life span can reach 25-30 years with sexual maturity before 6 years (Gori et al., 2007; Weinberg and Weinberg, 1979).

The other gorgonian, *L. sarmentosa* also releases lecithotrophic larvae but in the late summer (September to October) and the PLD is unknown (Rossi and Gili, 2009). *L. sarmentosa* life span can reach 20 years with female sexual maturity within 2-3 years after settlement (Rossi and Gili, 2009).

*H. papillosa* is a simultaneous hermaphrodite which releases larvae in late summer (September to October; Becerro and Turon, 1992), presumably lecithotrophic. The PLD of *H. papillosa* larvae is unknown but PLD shorter than 12 hours has been consistently reported for other solitary ascidian species (Ayre et al., 1997). We did not find any data about the age at sexual maturity and the life span of *H. papillosa*. However, the ascidians are considered as highly invasive, particularly because of their rapid growth and early sexual maturity (Zhan et al., 2015), with some species such as C*iona intestionalis* complex and *Ciona savigniy*, reaching sexual maturity at the age of 2 months (Zhan et al., 2015) and continuous spawning (Carver et al., 2003).

*P. fascialis* displays both sexual and asexual reproduction. During sexual reproduction, most bryozoans release lecithotrophic larvae which settle after a few minutes or a few hours, rarely beyond several days (Keough, 1983). *P. fascialis* larval release has been inferred to happen in June based on recruitment observations (Cocito et al., 1998a). Asexual reproduction happens by colony fragmentation or budding extension (Cocito et al., 1998b). Individual life span is estimated to be about 10 years with early sexual maturity after 2 years (Cocito et al., 1998b).

### Statistical analysis

We examined to which extent ARs colonization measured by the presence/absence of the five species and on the dissimilarity between their co-occurrence assemblages is affected by the AR shape, location and timing of deployment (age). Due to the GoL AR deployment set up, the effect of some factors was tested only locally. We first verified that shape was not affecting colonization, by testing the effect of shape in one geographical sector where different shapes were deployed at same depth and time (VLR1/VLR2, Table 1: Supplementary Material). Second, the effect of factors age (2 levels: 1985 and the 2000s) and depth (2 levels, >20 m, <20 m depth) was tested within the AGD geographical sector only. Factor age was tested in 3 sites at <20 m (AGD1/ AGD2 & AGD3) and factor depth was tested in 4 sites deployed in 2009 (AGD2&AGD3 / AGD4&AGD5). Third, factor location was tested both at regional and local levels (geographical sector being the regional factor, site being the local factor). To avoid any confounding effect due to age or depth, the regional factor vs the local factor were tested on the 5 geographical sectors where AR were deployed during the 2000s immersion phase and at <20 m depth only (AGM, AGD, VLR, GRU, LEU: 5×2 levels).

A Jaccard similarity matrix was built on presence/absence data across all pairwise sampling units used in each test (Table 2: supplementary material). Two multivariate analyses were performed. Non-parametric multivariate analysis of variance with permutation was applied to test for the effects of shape, age, depth and geographical sector on species assemblages (NP-manova: Anderson, 2001; Zar, 1999). Site was considered as a random factor, nested either in depth, age or the geographical sector. Another multivariate analysis was performed to cluster most similar species assemblages in the sector of AGD (SIMPROF: Clarke et al., 2008) (Table 2: supplementary material).

**Table 2.**
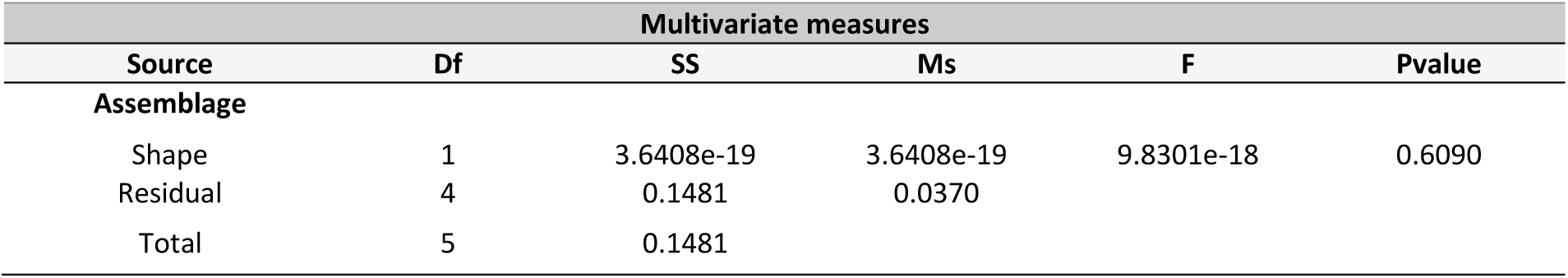
Results of multivariate NP-Manova testing the effect of the shape between steel cage and pipe on the presence/absence assemblage. Sites tested: VLR1 vs VLR2. Significant (P<0.05) values in bold.

| Multivariate measures |  |  |  |  |  |
| --- | --- | --- | --- | --- | --- |
| Source | Df | SS | Ms | F | Pvalue |
| Assemblage |  |  |  |  |  |
| Shape | 1 | 3.6408e-19 | 3.6408e-19 | 9.8301e-18 | 0.6090 |
| Residual | 4 | 0.1481 | 0.0370 |  |  |
| Total | 5 | 0.1481 |  |  |  |

When significant differences between the five species co-occurrence assemblages were detected for a factor, a non-parametric univariate analysis (Kruskall-Wallis) was performed for each species independently to detect the species driving the difference (Table 2: supplementary material). Fisher posthoc test was used to identify the site where the difference arose. A same p-value of 0.05 was taken for detecting significant differences. Analyses were performed with Matlab software using the Fathom package for multivariate analyses (Jones, 2014) and the Matlab statistics toolbox for univariate analyses.

## Results

### Artificial reefs colonization by the five target species at regional scale

Among the five target species, *S. spallanzanii* was the only one whose presence was recorded in all the sampling units and geographical sectors (Figure 3). In only one out of the 16 sites, it was the only species detected. *H. papillosa* and *L. sarmentosa* were detected in five of the six geographical sectors (not present in GRU and AGM respectively). *P. fascialis* and *E. singularis* were detected in 3 of the 6 geographical sectors (CST, LEU, and AGD for *P. fascialis* and AGM, AGD and LEU for *E. singularis*). Finally, *E. singularis* was the least frequently observed species, being detected in only 5 out of the 16 sites. In all geographical sectors, at least three of the five target species were detected, except GRU where only two of the five species were detected. Assemblages of two species were found in two sites out of the 16 (sectors AGD and GRU), assemblages of three species were found in 7 sites, assemblages of four species were found in 2 sites and assemblages of five species were found in 3 sites.

**Figure 3.**
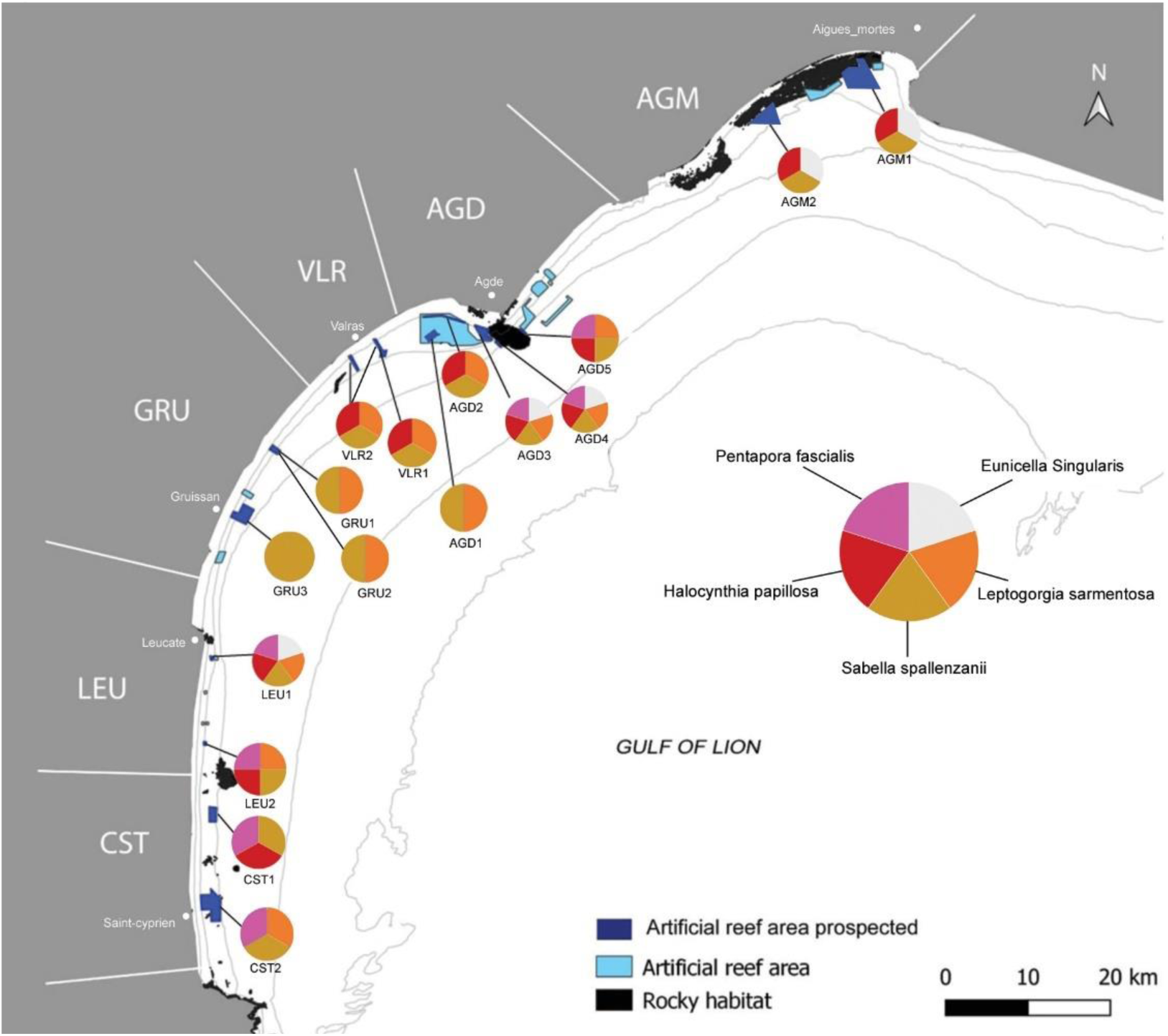
Map showing the five species co-occurence assemblages inventoried on ARs in the 16 sites (AGM1, AGM2, AGD1, AGD2, AGD3, AGD4, AGD5, VLR1, VLR2, GRU1, GRU2, GRU3, LEU1, LEU2, CST1, CST2) in the Gulf of Lion.

### Shape effect on five target species at local scale

No difference in species assemblages was found in VLR between pipe and steel cage (NP-MANOVA, F=9.08 e-19, P>0.05; Table 2).

Similarly, in the geographical sector GRU, the same assemblages were detected on sites differing by ARs shape (pipes and a heap of telegraphical poles) in the same location (GRU1 and GRU2). Conversely, different assemblages were detected between two sites (GRU2 and GRU3) separated by 9 km although ARs shape was the same (heaps of telegraphical poles). Due to the absence of replication in unit samplings, it was not possible to perform a statistical test on the effect of shape in this geographical sector.

### Age and depth effects on five target species at local scale

Despite all five target species being detected on ARs in the AGD sector, assemblage composition among sites differed (Figure 3). Clustering of sampling units within the 5 sites (AGD1, AGD2, AGD3, AGD4, AGD5) in AGD identified 2 clusters (SIMPROF: P<0.05; Figure 4). The two sites (AGD2 and AGD3) with same age (2009), depth range (less than 20 m) and reef shape (pipe) were attributed to different clusters. In fact, one cluster grouped ARs of different ages at a same depth (1985 in site AGD1 and 2009 in site AGD2) while the other cluster grouped ARs of the same age but at different depths (less than 20 m in site AGD3 and more than 20 m in sites AGD4 and AGD5). In both clusters, different AR shapes were found (steel cage and pipes in one cluster, pipes, Comin and Bonna in the other cluster) (Figure 4). The geographic distance between the two clusters (AGD4-AGD5) and (AGD3-AGD4-AGD5) was 7.5 km and the median value of the intra-cluster geographic distance was 3 km.

**Figure 4.**
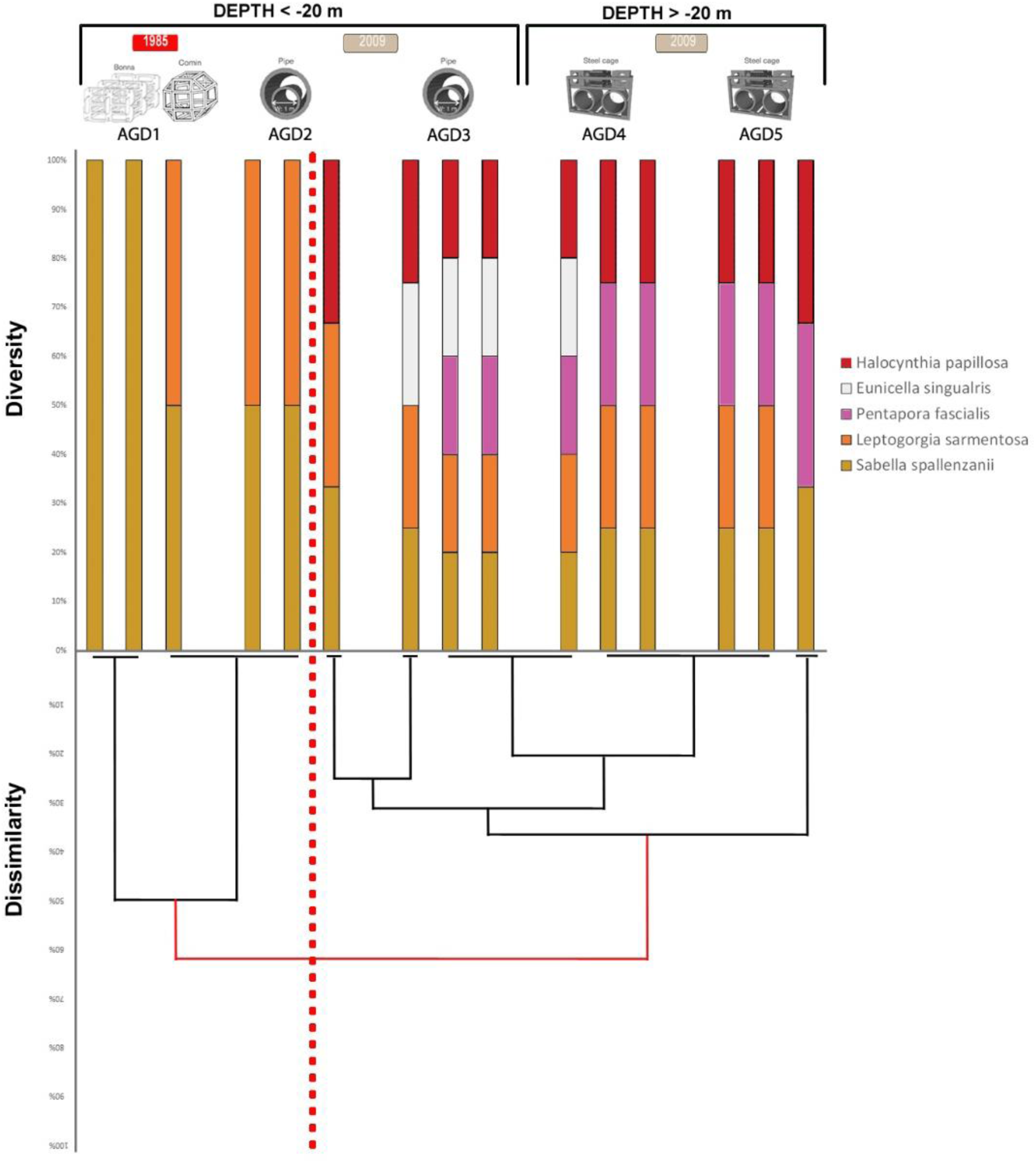
Diversity and dissimilarity. Upper part: diversity of species assemblages in the sampling units of the five sites (AGD1, AGD2, AGD3, AGD4, AGD5) in AGD sector together with the type of AR, depth and years of deployment. Lower part: dendrogram obtained by group average clustering based on the Jaccard dissimilarity index using the presence /absence of species (P=0.04 at 62% of dissimilarity). The red dotted line separates the two clusters identified by the analysis.

Multivariate analysis of variance confirmed that neither age (NP-MANOVA, F=1.43 P<0.05; Table 3) nor depth (NP-MANOVA, F=1.37 P<0.05; Table 4) explained site differences in the five species assemblages found on ARs in AGD (P<0.05; Tables 3 and 4).

**Table 3.** Results of multivariate NP-Manova testing the interactive effects of the age (of deployment) and site (nested in Age) on the presence/absence assemblage. Sites tested: AGD1 vs (AGD2 vs AGD3). Significant (P<0.05) values in bold.

| Multivariate measures |  |  |  |  |  |
| --- | --- | --- | --- | --- | --- |
| Source | Df | SS | Ms | F | Pvalue |
| **Assemblage** |  |  |  |  |  |
| Age | 1 | 0.4825 | 0.4825 | 1.4330 | 0.3330 |
| Site (Age) | 1 | 0.3367 | 0.3367 | 7.5551 | **0.0130** |
| Residual | 6 | 0.2674 | 0.0446 |  |  |
| Total | 8 | 1.0866 |  |  |  |

**Table 4.** Results of multivariate NP-Manova testing the interactive effects of the depth (of deployment) and site (nested in depth) on the presence/absence assemblage. Sites tested: (AGD2 vs AGD3) vs (AGD4 vs AGD5). Significant (P<0.05) values in bold.

| Multivariate measures |  |  |  |  |  |
| --- | --- | --- | --- | --- | --- |
| Source | Df | SS | Ms | F | Pvalue |
| **Assemblage** |  |  |  |  |  |
| Depth | 1 | 0.2491 | 0.2491 | 1.3715 | 0.3280 |
| Site(Depth) | 2 | 0.3633 | 0.1816 | 8.5969 | **0.001** |
| Residual | 8 | 0.1690 | 0.0211 |  |  |
| Total | 11 | 0.7816 |  |  |  |

Differences among the five sites in AGD were due to different ARs colonization by three species (Kruskall-Wallis: *E. singularis, P. fascialis, H. papillosa*; all P<0.05; Table 5). Site AGD3 differed from sites AGD5, AGD1 and AGD2 by the presence of *E. singularis*, sites AGD4 and AGD5 differed from sites AGD1 and AGD2 due to the presence of *P. fascialis* and the site AGD1 differed from sites AGD5, AGD3 and AGD4 by the presence of *H. papillosa* (Post-hoc tests, Table 3: supplementary material).

**Table 5.** Results of univariate Kruskall-Wallis testing the effect of location of deployment on the presence/absence by species. Sites tested: all sites in AGD. Significant (P<0.05) values in bold.

| Univariate measures |  |  |  |  |  |
| --- | --- | --- | --- | --- | --- |
| Source | Df | SS | Ms | Chi_sq | Pvalue |
| **Species** |  |  |  |  |  |
| Eunicella singularis | 4 | 127.5 | 31.875 | 10.82 | **0.0287** |
| Leptogorgia sarmentosa | 4 | 60 | 15 | 6.22 | 0.1832 |
| Pentapora fascialis | 4 | 172.5 | 43.125 | 11.5 | **0.0215** |
| Sabella spallanzanii | 4 | 0 | 0 | nan | nan |
| Halocynthia papillosa | 4 | 150 | 37.5 | 11.2 | **0.0244** |

### Geographical effect on five target species at local and regional scales

The five species co-occurrence assemblages on ARs deployed in the same period and at the same depth were significantly different at both regional and local scales (NP-MANOVA): geographical sector F=5.09 P<0.05; site (geographical sector) F=2.78 P<0.05; Table 6). These differences were due to different colonization of ARs by three of the five species, *E. singularis, L. sarmentosa* and *P. fascialis*. For the latter two species, regional differences (Kruskall-Wallis: geographical sector P=0.0002) were more significant than local differences (Kruskall-Wallis: site P=0.001 for *L. sarmentosa* and P=0.005 for *P. fascialis*; Table 7). Both species were not detected in the north of the GoL (AGM). In contrast, for *E. singularis*, local differences (Kruskall-Wallis: site P=0.017) were more significant than regional ones (geographical sector P=0.036; Table 7), the species being detected in geographical sectors in the north, center and south of GoL.

**Table 6.** Results of multivariate NP-Manova testing the interactive effects of geographical sector and site (nested in geographical sector) on the presence/absence assemblage. Sites tested: (AGM1 vs AGM2) vs (AGD2 vs AGD3) vs (VLR1 vs VLR2) vs (GRU1 vs GRU2 vs GRU3) vs (LEU1 vs LEU2). Significant (P<0.05) values in bold.

| Multivariate measures |  |  |  |  |  |
| --- | --- | --- | --- | --- | --- |
| Source | Df | SS | Ms | F | Pvalue |
| **Assemblage** |  |  |  |  |  |
| Geographical sector | 4 | 2.1671 | 0.5418 | 5.0994 | **0.005** |
| Site (Geographical sector) | 6 | 0.6375 | 0.1062 | 2.7812 | **0.007** |
| Residual | 18 | 0.6876 | 0.0382 |  |  |
| Total | 28 | 3.4921 |  |  |  |

**Table 7.** Results of univariate Kruskall-Wallis testing the effects of geographical sector and site on the presence/absence by species. Sites tested: AGM1 vs AGM2 vs AGD2 vs AGD3 vs VLR1 vs VLR2 vs GRU1 vs GRU2 vs GRU3 vs LEU1 vs LEU2. Significant (P<0.05) values in bold.

| Univariate measures |  |  |  |  |  |
| --- | --- | --- | --- | --- | --- |
| Source | Df | SS | Ms | Chi_sq | Pvalue |
| **Eunicella singularis** |  |  |  |  |  |
| Geographical sector | 4 | 447.08 | 111.77 | 10.28 | **0.036** |
| Site | 9 | 937.66 | 93.76 | 21.56 | **0.0175** |
| **Leptogorgia sarmentosa** |  |  |  |  |  |
| Geographical sector | 4 | 965.7 | 241.42 | 22.2 | **0.0002** |
| Site | 9 | 1116.5 | 11.65 | 28 | **0.0018** |
| **Pentapora fascialis** |  |  |  |  |  |
| Geographical sector | 4 | 937.66 | 234.41 | 21.56 | **0.0002** |
| Site | 9 | 1077.83 | 107.78 | 24.78 | **0.0058** |
| **Sabella spallanzanii** |  |  |  |  |  |
| Geographical sector | 4 | 0 | 0 | nan | nan |
| Site | 9 | 0 | 0 | nan | nan |
| **Halocynthia papillosa** |  |  |  |  |  |
| Geographical sector | 4 | 323.59 | 80.89 | 6.01 | 0.1985 |
| Site | 9 | 681.5 | 68.15 | 12.53 | 0.2509 |

## Discussion

The resilience of marine natural biodiversity, including benthic invertebrates, is essential to ensure the sustainability of ecosystem functions. Increasing habitat surface is key to support the natural biodiversity resilience, which in the case of the rocky habitat can be achieved by effectively integrated ARs. The effective integration of ARs into the rocky habitat network starts with their colonization by species from the natural habitat. In the present study the five species selected from the natural rocky habitat of the GoL colonized differently the ARs spread along the GoL coastline ten years after their deployment. Locally, neither age, immersion, depth or shape of the ARs significantly affected colonization patterns.

Colonization of ARs are expected to evolve toward a stable state comparable to that of the natural environment, through the succession of opportunistic species (wide dispersal, high fertility, low tolerance of reduced resource levels, short life-spans, minimal dietary specialisation) followed by specialized species (limited dispersal, slow growth to a large size at maturity, delayed and limited reproduction, optimization to reduced resources and long life spans; Platt and Connell, 2003; Faurie et al., 2003). Monitoring of ARs short-term colonization (<3 years) have indeed shown a dominance of pioneer species (hydroids, serpulids, barnacles and bivalves), most of them having life history traits typical of opportunistic species (Fariñas-Franco et al., 2013; T. Glasby, 1999; Moura et al., 2007; Pamintuan and Ali, 1994; Ponti, 2015; Relini et al., 1994; Spagnolo et al., 2014; Toledo et al., 2020).

Long-term studies confirmed successions in ARs colonization (Burt et al., 2011; Nicoletti et al., 2007; Perkol-Finkel and Benayahu, 2005; Whomersley and Picken, 2003), but none have described saturation (Svane and Petersen, 2001). In the Tyrrhenian Sea, Nicolleti et al. (2007) described colonization in 5 distinct phases: (i) a first recruitment by pioneer species (hydroids, serpulids, barnacles and bivalves) during the firsts months after immersion, followed by phases of (ii) cover dominance, (iii) regression and (iv) absence of *Mytilus galloprovincialis* for more than 10 years. The installation of diverse bio-builders bryozoans characteristic of the natural environment was recorded after 20 years only (v). Our study shows that biobuilder engineering species such as bryozoans (*P. fascialis*) and gorgonians (*E. singularis, L. sarmentosa*) colonized ARs as early as 10 years after their deployment, without significant difference between 10 years and 35 years old ARs. However, the presence of *S. spallanzanii* described as an opportunistic species (sexual precocity, various reproduction modes, rapid growth, short lived; Giangrande et al., 2000) on all ARs independently of their age of deployment suggests that ARs did not yet reach a stable state comparable to the natural environment. Thus, the presence of bio-builders is not a sufficient indicator of the ARs naturalization to the local biodiversity.

The GoL’s ARs being located in the sandy coastal zone are likely regularly disturbed by sediment deposits due to river delivery or/and their resuspension by either trawling activities or the mechanical action of the swell (Dufois et al., 2014; Durrieu de Madron et al., 2005; Ulses et al., 2008). Testing the impact of swell and sediment deposit on ARs requires exploring the colonization of ARs along a gradient of depth and distance from the coast (Van der Stap et al., 2016). However, current ARs deployment in the GoL ranged from 15 to 30 m depth and within 3 miles from the coast and did not allow testing for differential effect of sediment disturbances as swell impact occurs every year in this area (Guizien, 2009). Testing the impact of sediment disturbance on ARs colonization would require exploring reefs deployed deeper than 50 m, such as the anchorages of the floating wind farm that will be placed in the GoL in the next future (https://infoefgl.fr/le-projet/le-parc/#). Light is also expected to be an important factor structuring benthic assemblages, along a depth gradient in natural and artificial environments (Glasby, 1999a; Glasby, 1999b; Svane and Petersen, 2001). Absence of depth effect in the present study, although in the GoL light intensity strongly attenuates within the upper 30 m of the water column (Durrieu de Madron et al., 2011), was potentially a bias due to the five species selected in the present study whose distributions are not strongly structured by light intensity.

Another factor which has been shown to drive the intensity of ARs colonization is structural complexity (see Bohnsack and Sutherland, 1985 for a review). Nevertheless, there is no consensus about the relationship between complexity and subtidal benthic invertebrates abundance, due to potential bias in controlling the surface and scale in ARs of different complexity (Rouse et al., 2019; Strain et al., 2018).

The similarity in the 5 species co-occurrence between different reef shapes at the same depth and of the same age found in the present study suggests that shape is less important than the geographical location in AR colonization by benthic invertebrates when controlling the colonized surface. However, shape is an imprecise measure of structural complexity. The latter is rarely assessed quantitatively and can be described by different metrics which may be similar for apparently different shapes (such as steel cages and pipes; Riera, 2020).

Benthic invertebrate assemblages result from complex processes that operate at multiple spatial and temporal scales (Smale, 2012). At the regional scale, larval availability can become a major factor explaining colonization success (Padron et al., 2018). Change in the composition of assemblages during the early colonization of artificial substrates by benthic invertebrates has been attributed to the availability and abundance of larvae during the seasons rather than a sequence of distinct succession (Bramanti et al., 2003; Turner and Todd, 1993). The larval behaviour (buoyancy and motility) and the characteristics of the biological cycle of the species (spawning timing and PLD) can play a key role in determining the dispersal distance (Todd, 1998), and consequently the possibility to reach habitat suitable for settlement. Dispersion distance is generally correlated with PLD, thus a species with a long PLD is supposed to colonize habitats further away than species with a shorter PLD (Shanks, 2009). In this study, the five species were chosen among different phyla known for their contrasting planktonic durations, swimming abilities and larval dispersal periods, although these larval traits are only known accurately for *E. singularis* (Guizien et al., 2020; Zelli et al., 2020). *P. fascialis* and *H. papillosa*, the two species with short PLD (<24h and <12h, respectively) colonized ARs located close to the natural habitat where they are present (<4.8 km and <10 km, respectively). The coastal circulation of the GoL allows such dispersal distance over periods of a few days (Guizien et al., 2012). *S. spallanzanii*, which has a PLD larger than 3 weeks, colonized all the inventoried ARs, in line with a dispersal distance of 40 km after 3 weeks (Guizien et al., 2012). In contrast, *E. singularis* did not colonize all ARs within geographical sectors of 30 km width where the species is present in the natural habitat, although a 2 week PLD was expected to enable such dispersal (Padron et al., 2018). The other gorgonian species, *L. sarmentosa* colonized more ARs located within distances of less than 30 km from its natural habitat than *E. singularis* while the PLD of the two species are presumably the same. This suggests that other factors influence the colonization failure of *E. singularis*.

Reproductive traits are another key to the success in colonising new settings (Stearns, 2000). In this regard, *E. singularis* colonization potential could be limited by its low fecundity (∼25-40 larvae.cm^-1^ of colony branch; Ribes et al., 2007; Theodor, 1967) compared to that of *L. sarmentosa* (∼75 larvae.cm^-1^ of colony branch; Rossi et al., 2011; Rossi and Gili, 2009). The wide colonization of ARs by *S. spallanzanii* is in line with its reproductive traits typical of opportunistic species (early sexual maturity, high fecundity with more than 50 000 eggs per female, Currie et al., 2000, a fertilization close to 100%, Giangrande et al., 2000). Since arriving in the Pacific Ocean, *S. spallanzanii* has been declared one of the ten priority pest species in the marine environment by the Australian authorities and classified as an invasive species (Hayes et al., 2005). Similarly to *S. spallanzanii, H. papillosa* colonized nearly all ARs located within its 10 km dispersal distance from the natural habitat. Within the ascidian class, a wide disparity in species fecundity has been reported (Pandian, 2018). This suggests *H. papillosa* reproductive traits would be close to those of invasive ascidians (Zhan et al., 2015).

Ultimately, understanding ARs colonization requires a precise mapping of source populations in the natural environment. To this respect, the abundance of *S. spallanzan*i on ARs is surprising, as the species is not abundant in the natural rocky habitat of the GoL. For this species, other sources of larval supply than natural settings should be considered, such as the numerous ports and marinas along the coast of the GoL, as *S. spallanzanii* is very tolerant to environmental conditions (Currie et al., 2000). In this case of intense colonization by an endemic benthic invertebrate species, ARs apparently extended its metapopulation, acting as stepping stones for further larval dispersal beyond its natural current colonization limits (Bishop et al., 2017; Wang et al., 2020).

In the GoL, the 14 500 m^3^ of ARs deployed 30 years ago are now decommissioned and the relevance of their removal is currently debated. Connectivity between natural populations has been shown to support species resilience after disturbances in fragmented habitat, and could be extended to ARs (Fahrig, 2003). However, ARs may also facilitate the spread of non-indigenous species introduced with maritime traffic in ports (Glasby et al., 2007).

The present study advocates accounting for the geographical arrangement in planning ARs deployment to extend the habitat of hard bottom benthic invertebrate natural populations.

## Data accessibility

Data are available online https://doi.org/10.5281/zenodo.5567023

## Supplementary material

Script and codes are available online: https://doi.org/10.1101/2021.10.08.463669

**Supplementary Table 1:** Stratified sampling design descriptors. Sector, site, latitude and longitude of the sampling unit centroid, depth of immersion, year of deployment, type, number and surface of AR inventoried per sampling unit.

**Supplementary Table 2:** Factors and sites used for multivariate and univariate testing.

**Supplementary Table 3:** Results of post-hoc test of univariate Kruskall-Wallis testing the effect of location of deployment on the presence/absence for *E. singularis, L. sarmentosa and H. papillosa*. Sites tested: all sites in AGD. Significant (P<0.05) values in bold.

## Acknowledgements

This work was funded by the Agence de l’Eau Rhône-Méditerranée-Corse under project ICONE - Impacts actuels et potentiels de la CONnectivité Ecologique ajoutée par les récifs artificiels sur la biodiversité fixée des substrats durs du Golfe du Lion (PI, K. Guizien, AAP 2016). The authors gratefully acknowledge the helpful assistance during sampling of the staff of the Aire Marine Protégée Agathoise. Version 4 of this preprint has been peer-reviewed and recommended by Peer Community In Ecology (https://doi.org/10.24072/pci.ecology.100093)

## Authors contribution

SB and KG conceived the study, SB carried out sampling and statistical analysis. All contributed to manuscript writing.

