## Supplementary Material for "Artificial reefs geographical location matters more than shape, age and depth for sessile invertebrate colonization in the Gulf of Lion (NorthWestern Mediterranean Sea)"

**ABSTRACT**

Artificial reefs (ARs) have been used to support fishing activities. Sessile invertebrates are essential components of trophic networks within ARs, supporting fish productivity. However, colonization by sessile invertebrates is possible only after effective larval dispersal from source populations, usually in natural habitat. While most studies focused on short term colonization by pioneer species, we propose to test the relevance of geographic location, shape, age and depth of immersion on the ARs long term colonization by species found in natural stable communities in the Gulf of Lion. We recorded the presence of five sessile invertebrates species, with contrasting life history traits and regional distribution in the natural rocky habitat, on ARs with different shapes deployed during two immersion time periods (1985 and the 2000s) and in two depth ranges (<20m and >20m). At the local level (~5kms), neither shape, depth nor immersion duration differentiated ARs assemblages. At the regional scale (>30kms), colonization patterns differed between species, resulting in diverse assemblages. This study highlights the primacy of geographical positioning over shape, immersion duration and depth in ARs colonization, suggesting it should be accounted for in maritime spatial planning.

**KEY WORDS:** ARTIFICIAL REEF, BENTHIC INVERTEBRATES, SHAPE, IMMERSION DEPTH , AGE, GEOGRAPHICAL LOCATION, LIFE TRAITS

Table 1 : Stratified sampling design descriptors. Sector, site, latitude and longitude of the sampling unit centroid, depth of immersion, year of deployment, type, number and surface of AR inventoried per sampling unit.

**Table 1 :**

| Sector / Surface of rocky natural habitat in the sector (hectare) | Site / Total sampled surface (m2) | Latitude (WGS 84) | Longitude (WGS 84) | Depth (m) | Year of deployment | AR shape | Number of ARs inventoried | Sampling unit surface (m²) |
|---|---|---|---|---|---|---|---|---|
| Golfe d'Aigues Mortes (AGM) / 3123 | AGM1 / 720 | 43.5242 | 4.03964 | 16 | 2006 | Pyramid | 1 | 240 |
| | | | | | | | 1 | 240 |
| | | | | | | | 1 | 240 |
| | AGM2 / 720 | 43.4851 | 3.9042 | 16 | 2006 | Pyramid | 1 | 240 |
| | | | | | | | 1 | 240 |
| | | | | | | | 1 | 240 |
| Agde (AGD)/ 1168 | AGD1 / 434 | 43.2465 | 3.4052 | 17 | 1985 | Bonna | 1 | 160 |
| | | | | | | | 1 | 160 |
| | | | | | 1985 | Comin | 2 | 114 |
| | AGD4 / 405 | 43.2400 | 3.4969 | 28 | 2009 | Steel basket | 1 | 135 |
| | | | | | | | 1 | 135 |
| | | | | | | | 1 | 135 |
| | AGD5 / 405 | 43.2535 | 3.5356 | 27 | 2009 | Steel basket | 1 | 135 |
| | | | | | | | 1 | 135 |
| | | | | | | | 1 | 135 |
| | AGD2 / 306 | 43.2689 | 3.4114 | 10 | 2009 | Pipe | 3 | 102 |
| | | | | | | | 3 | 102 |

| | | | | | | | | |
|---|---|---|---|---|---|---|---|---|
| | | | | | | | 3 | 102 |
| | AGD3 / 544 | 43.2551 | 3.47375 | 10 | 2009 | Pipe | 4 | 136 |
| | | | | | | | 4 | 136 |
| | | | | | | | 4 | 136 |
| Valras (VLR) / 76 | VLR1 / 405 | 43.2299 | 3.3264 | 20 | 2006 | Steel basket | 1 | 135 |
| | | | | | | | 1 | 135 |
| | | | | | | | 1 | 135 |
| | VLR2 / 374 | 43.2330 | 3.3004 | 12 | 2006 | Pipe | 4 | 136 |
| | | | | | | | 4 | 136 |
| | | | | | | | 3 | 102 |
| Gruissan ( GRU) /0 | GRU1 / 306 | 43.1253 | 3.1615 | 14 | 2004 | Pipe | 3 | 102 |
| | | | | | | | 3 | 102 |
| | | | | | | | 3 | 102 |
| | GRU2 / 459 | 43.1253 | 3.1615 | 14 | 2004 | heaps of telegraph poles | 1 | 459 |
| | GRU3 / 459 | 43.055 | 3.1033 | 12 | 2002 | heaps of telegraph poles | 1 | 459 |
| Leucate Barcarès (LEU) / 50 | LEU1 / 445 | 42.8941 | 3.0696 | 17 | 2004 | Concrete box | 2 | 178 |
| | | | | | | | 2 | 178 |
| | | | | | | | 1 | 89 |
| | LEU2 / 445 | 42.8237 | 3.0577 | 18 | 2004 | Concrete box | 2 | 178 |
| | | | | | | | 2 | 178 |
| | | | | | | | 1 | 89 |

| | | | | | | | | |
|---|---|---|---|---|---|---|---|---|
| St cyprien-Canet (CST) / 1249 (756 + 493) | CST1 / 434 | 42.7243 | 3.0723 | 29 | 1985 | Bonna | 1 | 160 |
| | | | | | | | 1 | 160 |
| | | | | | | Comin | 2 | 114 |
| | CST2 / 434 | 42.6253 | 3.0707 | 30 | 1985 | Bonna | 1 | 160 |
| | | | | | | | 1 | 160 |
| | | | | | | Comin | 2 | 114 |

**Table 2 :** Factors and sites used for multivariate and univariate testing.

| **Factors and sites testing** | | |
|---|---|---|
| Multivariate measures | | |
| **Factor1** | **Factor2** | **Tested sites** |
| Shape | | VLR1 vs VLR2 |
| Age | Site | AGD1 vs (AGD2 vs AGD3) |
| Depth | Site | (AGD2 vs AGD3) vs (AGD4 vsAGD5) |
| Geographical sector | Site | (AGM1 vs AGM2) vs (AGD2 vs AGD3) vs (VLR1 vs VLR2) vs (GRU1 vs GRU2 vs GRU3) vs (LEU1 vs LEU2) |
| Univariate measures | | |
| Location | | AGD1 vs AGD2 vs AGD3 vs AGD4 vs AGD5 |
| Geographical sector | | AGM AGD VLR GRU LEU |
| Site | | AGM1 vs AGM2 vs AGD2 vs AGD3 vs VLR1 vs VLR2 vs GRU1 vs GRU2 vs GRU3 vs LEU1 vs LEU2 |

**Table 3 :** Results of post-hoc test of univariate Kruskall_wallis testing the effect of location of deployment on the presence/absence for *E. singularis, L. sarmentosa, H. papillosa*. Sites tested: all site in AGD. Significant (P<0.05) values in bold.

| Univariate measures | | | | |
|---|---|---|---|---|
| **Post-hoc** | | | | |
| **SITE** | **SITE** | Mean difference | Std.error | **Pvalue** |
| *Eunicella singularis* | | | | |
| AGD5 | AGD4 | -7,99 | 2,99 | 0,37 |
| **AGD5** | **AGD3** | **-12,99** | **-2,01** | **0,01** |
| AGD5 | AGD2 | -5,49 | 5,49 | 1,00 |
| AGD5 | AGD1 | -5,49 | 5,49 | 1,00 |
| AGD4 | AGD3 | -10,49 | 0,49 | 0,07 |
| AGD4 | AGD2 | -2,99 | 7,99 | 0,37 |
| AGD4 | AGD1 | -2,99 | 7,99 | 0,37 |
| **AGD3** | **AGD2** | **2,01** | **12,99** | **0,01** |
| **AGD3** | **AGD1** | **2,01** | **12,99** | **0,01** |
| *Pentapora fascialis* | | | | |
| AGD5 | AGD4 | **-7,99** | 6,20 | 1,00 |
| AGD5 | AGD3 | **-12,99** | 8,70 | 0,43 |
| **AGD5** | **AGD2** | **-5,49** | **13,70** | **0,02** |
| **AGD5** | **AGD1** | **-5,49** | **13,70** | **0,02** |
| AGD4 | AGD3 | **-10,49** | 8,70 | 0,43 |
| **AGD4** | **AGD2** | **-2,99** | **13,70** | **0,02** |
| **AGD4** | **AGD1** | **-2,99** | **13,70** | **0,02** |
| AGD3 | AGD2 | **2,01** | 11,20 | 0,11 |
| AGD3 | AGD1 | **2,01** | 11,20 | 0,11 |
| AGD2 | AGD1 | **-5,49** | 6,20 | 1,00 |
| *Halocynthia papillosa* | | | | |
| AGD5 | AGD4 | -5,86 | 5,86 | 1,00 |
| AGD5 | AGD3 | -5,86 | 5,86 | 1,00 |
| AGD5 | AGD2 | -0,86 | 10,86 | 0,09 |
| **AGD5** | **AGD1** | **1,64** | **13,36** | **0,01** |
| AGD4 | AGD3 | -5,86 | 5,86 | 1,00 |
| AGD4 | AGD2 | -0,86 | 10,86 | 0,09 |
| **AGD4** | **AGD1** | **1,64** | **13,36** | **0,01** |
| AGD3 | AGD2 | -0,86 | 10,86 | 0,09 |
| **AGD3** | **AGD1** | **1,64** | **13,36** | **0,01** |
| AGD2 | AGD1 | -3,36 | 8,36 | 0,40 |
| AGD5 | AGD4 | -5,86 | 5,86 | 1,00 |
| AGD5 | AGD3 | -5,86 | 5,86 | 1,00 |
